# Integrate Heterogeneous NGS and TGS Data to Boost Genome-free Transcriptome Research

**DOI:** 10.1101/2020.05.27.117796

**Authors:** Yangmei Qin, Zhe Lin, Dan Shi, Mindong Zhong, Te An, Linshan Chen, Yiquan Wang, Fan Lin, Guang Li, Zhi-Liang Ji

## Abstract

It is a long-term challenge to undertake reliable transcriptomic research under different circumstances of genome availability. Here, we newly developed a genome-free computational method to aid accurate transcriptome assembly, using the amphioxus as the example. Via integrating ten next generation sequencing (NGS) transcriptome datasets and one third-generation sequencing (TGS) dataset, we built a sequence library of non-redundant expressed transcripts for the amphioxus. The library consisted of overall 91,915 distinct transcripts, 51,549 protein-coding transcripts, and 16,923 novel extragenic transcripts. This substantially improved current amphioxus genome annotation by expanding the distinct gene number from 21,954 to 38,777. We consolidated the library significantly outperformed the genome, as well as *de novo* method, in transcriptome assembly from multiple aspects. For convenience, we curated the Integrative Transcript Library database of the amphioxus (http://www.bio-add.org/InTrans/). In summary, this work provides a practical solution for most organisms to alleviate the heavy dependence on good quality genome in transcriptome research. It also ensures the amphioxus transcriptome research grounding on reliable data.

## Introduction

Transcriptome carries essential messages in depicting global gene expression landscape in biological events. Since last decade, next generation sequencing (NGS) has taken the leading position in large-scale gene expression determination under different philological conditions. Accompanying with the wide application of NGS, both reference-based and *de novo* algorithms were raised to interpret sequencing data. Most of the reference-based transcriptome assembly/annotation pipelines like TopHat (Trapnell, Pachter, & Salzberg)+Cufflinks(Cole Trapnell et al.) and HISAT2(Kim, Langmead, & Salzberg) are powerful to detect well-annotated genes consistently; however, they heavily rely on high quality genome (Moreton, Izquierdo, & Emes, 2015; Pertea, Kim, Pertea, Leek, & Salzberg, 2016) and are thus weak in representing RNA transcripts fully. In particular, they are unable to produce reliable transcriptome for organisms with poorly annotated genomes, and they do not work on no genome organisms. Alternatively, *de novo* methods like Trinity (Haas et al., 2013), IDBA (Peng, Leung, Yiu, & Chin, 2010), and Oases (Schulz, Zerbino, Vingron, & Birney, 2012) can reconstruct transcripts without genome reference. Their performances however largely depend on sequencing itself like library construction, sequencing length, and sequencing depth (Sohn & Nam, 2016). What’s more, complicated elements like repetitive sequences of varying length and copy number may also affect the quality of *de novo* assembly (Chaisson, Wilson, & Eichler, 2015). Therefore, the *de novo* assembled transcripts often consist of a number of fragmentary sequences. Another big setback for *de novo* approaches is that many transcripts are either falsely assembled, such as chimeric transcripts, or just discarded for too short sequence (Yang & Smith, 2013). A fraction of the discarded transcripts could be gene isoforms that may exert specific biological functions.

To neutralize the shortcomings of both reference-based and *de novo* methods, several approaches were developed to aid single transcriptome assembly. These approaches include the *de novo* and genome-guided assembly combination (Armero, Baudouin, Bocs, & This, 2017; X. Huang, Chen, & Armbruster, 2016; Orgeur et al., 2018), the multiple *de novo* approaches combination (Cerveau & Jackson, 2016; Nakasugi, Crowhurst, Bally, & Waterhouse, 2014; Rupp et al., 2014), the multiple-kmer usage (G. Chen, Yin, Wang, & Shi, 2011; Surget-Groba & Montoya-Burgos, 2010; Zhao et al., 2011), and the kmer-sequencing coverage depth combination (C. C. Chen, Lin, Chang, Chen, & Ho, 2012). These valuable efforts substantially improve the quality of single transcriptome assembly; however, they still highly rely on sequencing data itself and hardly overcome the defects of sequencing bias, fragmentary transcript, low expression transcript, and isoform.

Recent emergence of the third-generation sequencing (TGS) technologies bring the hope to solve the problem of accurate transcriptome assembly. In particular, the leading edges of long sequencing length and no PCR amplification largely alleviate the assembly difficulty in obtaining full-length transcripts (Sharon, Tilgner, Grubert, & Snyder, 2013). Unfortunately, the shortcomings of low throughput, inconsistent quantification, high error rate, low RNA capture rate, and high cost slow down the widespread use of TGS in biomedical research (Korlach, 2013; Lowe, Shirley, Bleackley, Dolan, & Shafee, 2017). Therefore, for the coming few years or a longer period, the NGS will still hold the leading place in large-scale transcriptome research, even for the rapid widespread of single cell sequencing recently.

For many under-studied animal and plant organisms, a number of gene products are just lost in the reference-based transcriptome (Bao, Xu, Jing, Meng, & Qin, 2015; Bouyioukos et al., 2013; Gan et al., 2011). Even for human and mouse that have the best genome annotation up-to-date, a portion of genes or isoforms cannot be fully detected via current computational or experimental approaches yet (Hu et al., 2015; Wu et al., 2014). Therefore, accurate transcriptome is still a pendent problem need to be solved at the first convenience.

In this study, we aim to develop a new computational method to aid accurate transcriptome reconstruction. For this purpose, we choose the amphioxus (lancelet) as an example to show how to maximally use heterogeneous data to assist accurate transcriptome assembly under the circumstance of having limited genome information. The amphioxus is a privileged species in phylogenic study of the invertebrate-to-vertebrate transition (Benito-Gutiérrez, 2011). In 2008, the US Joint Genome Institute (JGI) released the first amphioxus genome of *Branchiostoma floridae* (*B. floridae*), which is 522M base pair (bp) in size (N. H. Putnam et al., 2008). Thereafter, research groups in China and UK presented the genomes of *Branchiostoma belcheri* (*B. belcheri*) in 2014 (S. Huang et al., 2014) and amphioxus species *Branchiostoma lanceolatum* (*B. lanceolatum*) in 2018(Marlétaz et al., 2018), respectively (**Table S1**). Current assembly and annotation status of these three genomes is regretfully far away from perfectness. For instance, there exist many gaps in amphioxus genome that has not be filled up yet, which yields a number of scaffold fragments. Functional annotation of the amphioxus genomes is still rough since there is short of a good neighboring organism genome for reference that only 21,954 genes were annotated. Moreover, prior studies reported that amphioxus transcriptome only covered 41% of the genome predicted genes (Oulion, Bertrand, Belgacem, Le Petillon, & Escriva, 2012). These findings together implicate the difficulty of accurate transcriptome research if relying on the amphioxus genomes alone. It largely slows down the pace of the amphioxus on its road to the qualified animal model.

## Results

### Evaluation of the Integrative Transcript Library for the Amphioxus

#### Statistics of the transcript library

After quality control, ten RNA-seq datasets yielded about 613 million clean reads and 76.66 Giga (G) base pairs (bp) (**Table S2**). According to the released *B. floridae* genome of 522Mbp (Putnam et al, 2008), the average sequencing depth is about 73.4X. The TGS dataset yielded about 5.5 million subreads and 15.74Gbp in size (**Table S3**). In basis of these datasets, the computational workflow TransIntegrator produced a transcript library of 91,915 non-redundant sequences (**Figure 2**). This number of transcripts was about 4.2 times of gene number (21,954) annotated in the amphioxus genome (JGIv2.0). The integrity of all transcripts in the library were supported by multiple (>3) mapping reads. The majority (about 70.58%) of transcripts ranged 300bp to 2kbp in length, and the average length of the library transcripts was 1,667 bp (**Figure 2a**). This size is a bit larger than previous estimation of average amphioxus gene length of 1.4kbp (Nicholas H. Putnam et al.), indicating that the transcripts in the library were mostly well assembled. Out of the 91,915 sequences, about 12.7% (11,706) were protein-coding, 43.3% (39,843) were potentially protein-coding (ORFs without definite protein names), 4.8% (4,386) were potential long non-coding RNAs (lncRNAs), 0.4% (330) were other non-coding RNAs (ncRNAs), and the remaining 38.8% (35,650) were unannotated sequences.

**Figure 1.**
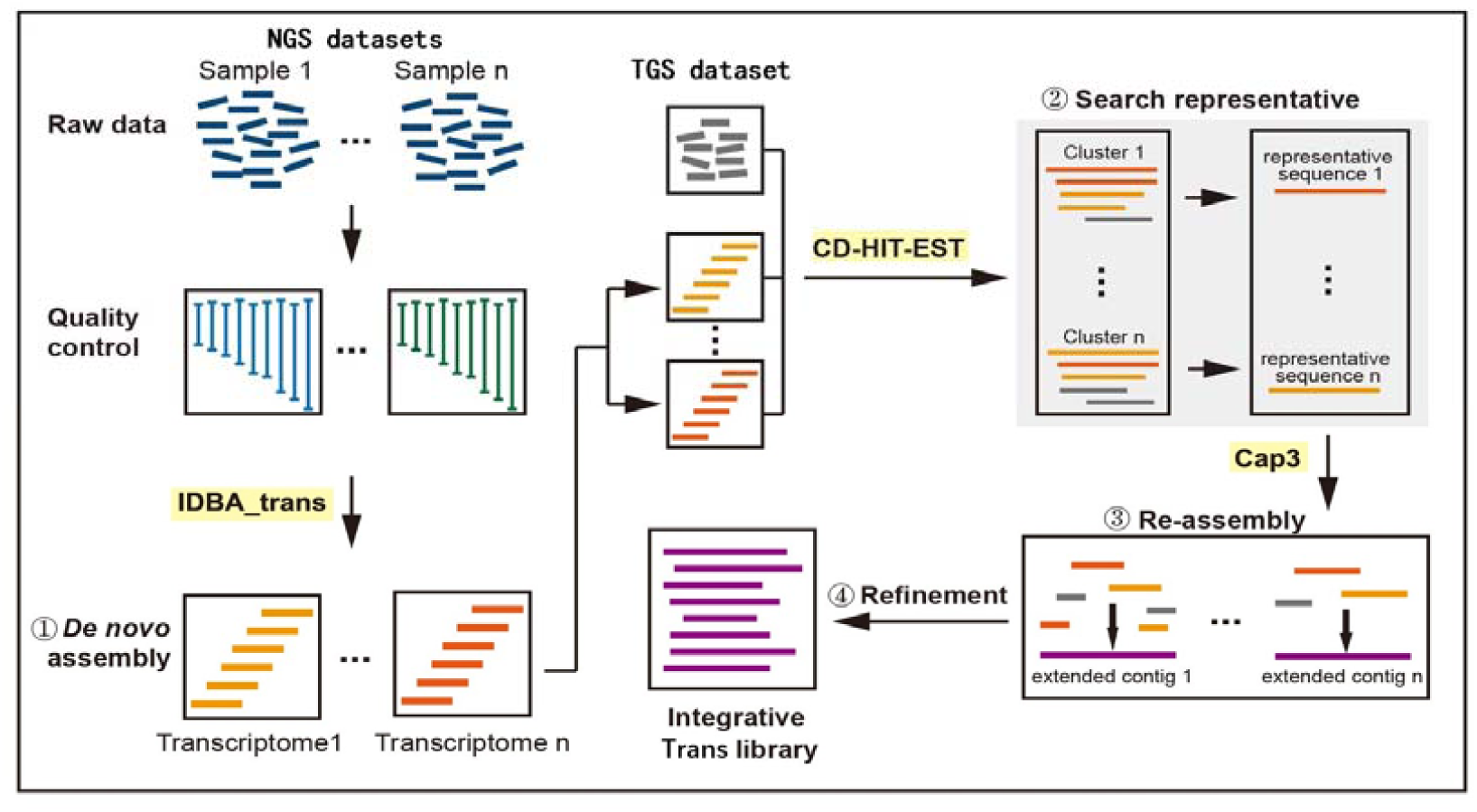
The computational workflow for construction of the integrative transcript library.

**Figure 2.**
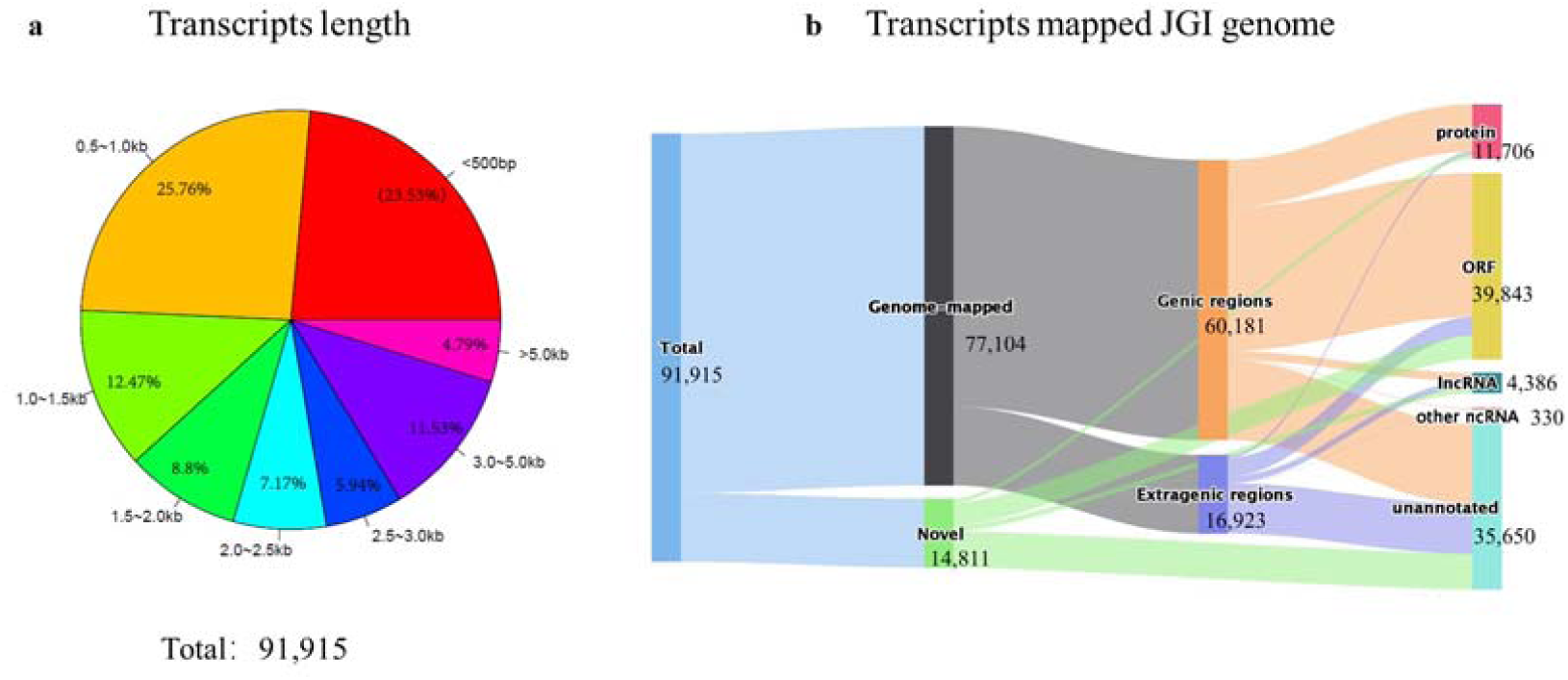
Validation and statistics of the integrative transcript library. (a) The length distribution of the library transcripts. (b) The functional annotation of the library transcripts. The library consists of overall 91,915 distinct transcripts, of which 11,706 are protein-coding, 39,843 are ORF-coding, 4,386 are lncRNAs, 330 are other ncRNAs, and the remaining 35,650 are unannotated sequences. Total 77,104 transcripts can be mapped to the amphioxus genome.

#### Comparison with the genome

We mapped the library transcripts against the amphioxus genome (JGIv2.0) by satisfying criteria of sequence identify >85% and sequence coverage >30%. The mapping identified the loci of about 83.9% (77,104 out of 91,915) of library sequences on the amphioxus genome. They hit about 97.0% (386 out of 398) of scaffolds and about 72.3% (15,867 out of 21,954) of genes. Besides, about 21.9% (16,923 out of 77,104) of the genome-mapped transcripts did not match the gene regions annotated by JGIv2.0 exactly. These extragenic transcripts included 4,628 potential protein-coding transcripts (600 protein-coding and 4,028 ORFs), 1,459 lncRNAs, 67 other ncRNAs, and 10,769 unannotated transcripts (**Figure 2b**). These extragenic transcripts complemented current amphioxus genome annotation by expanding the distinct gene number from 21,954 to 38,777.

In addition, the library also contained 14,811 “novel” transcripts that cannot find their loci on the genome. The novel transcripts included 5,888 protein-coding sequences, 1,112 lncRNAs, and 82 other ncRNAs (**Figure 2b**). Blasting these novel sequences against the NCBI RefSeq database denied the possibility of microbial contamination (sequence identify >90 and sequence coverage >90). These unmapped sequences may provide clues to fill up the gaps between genome scaffolds.

#### The heterogeneity and completeness of the library

The transcript library expanded in size along with the increase of RNA-seq datasets from one up to ten. However, the number of protein-coding transcripts reached the plateau of 49,698 after involving ten NGS transcriptomes (**Figure 3**). This amount is close to previous estimate of 50,817 proteins based on the genome annotation (N. H. Putnam et al., 2008). Besides, about 6,898 library transcripts can be mapped to the zebrafish genome, indicating the diversity of amphioxus genes have been fully represented. To the contrast, the number of non-coding transcripts continued to grow with the increase of datasets (**Figure 3a**), hinting their much more complicated and diverse roles during embryo development than what we thought previously.

**Figure 3.**
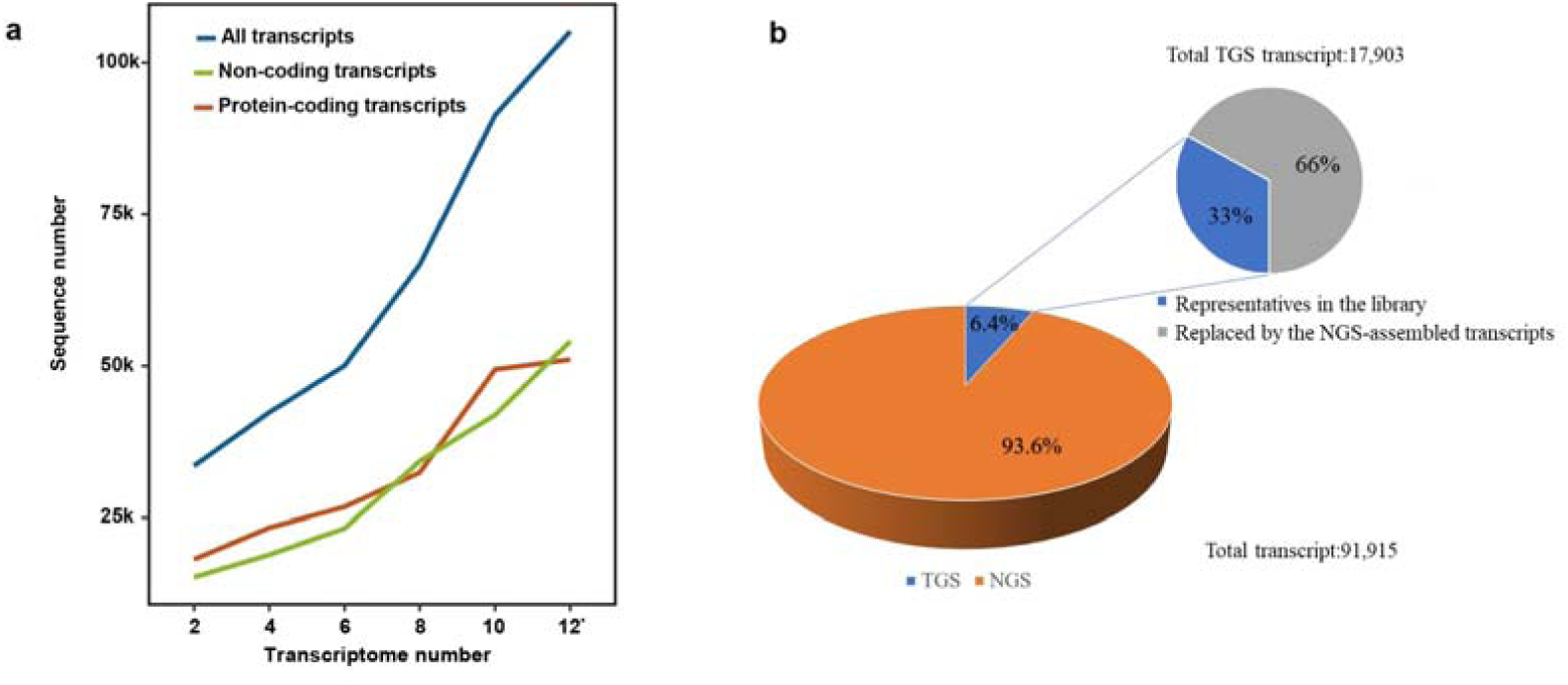
The scalability and heterogeneity of the integrative transcript library. (a) The scalability of the transcript library. The library grows with the increase of involved RNA-seq datasets, by both total transcripts and protein-coding transcripts. The protein-coding transcripts reach a plateau after involving 10 datasets. (b) The heterogeneous component of the transcript library. About 6.4% of library sequences are represent by transcripts derived from the TGS dataset.

Out of the overall 91,915 non-redundant transcripts in the library, the TGS dataset contributed 6.4% (5,900 transcripts), the NGS datasets contributed the remaining 93.6% (86,015 transcripts). Worthy of mention, involving TGS dataset did not lead to the substantial growth of the transcript library. About 2/3 of TGS-originated transcripts were replaced by the NGS-assembled transcripts due to the comparatively shorter length; and the remaining 1/3 (5900) transcripts however took placed of the corresponding NGS-assembled transcripts, which increased the protein-coding transcripts number from 49,698 to 51,549. The annotation and the distribution of the transcripts were illustrated by origin group in the **Figure 3b**.

### The Integrative Transcript Library Database for the Amphioxus

For user convenience, we curated an on-line database in basis of the integrative transcript library for the amphioxus (*B. floridae*), namely InTrans. The database was deployed on a Linux + Tomcat system architecture, and the interactive user-interfaces were designed using the JavaScript technology.

The InTrans database is now freely accessible at http://www.bio-add.org/InTrans/. The database allows transcript search via either browse or blast ways. The database implements the JBrowse genome browser (Buels, Yao, Diesh, Hayes, & Holmes, 2016) for better visualization of transcript locus on genome. In the genome browser, the transcripts are mapped back to the *B. floridae* genome (JGIv2.0), indicating their genomic loci and comparing with JGIv2.0 annotation. Clicking on the transcript will prompt the detailed information of the transcript, including the functional assignment (i.e. protein-coding, lncRNA, and so on), the exon-intron structure, and the corresponding sequences. Besides, the database also deploys blast function to enable search of transcript database via sequence alignment. The blast function allows input of both nucleotide and protein sequences in the *fasta* format via text form or uploaded file. The blasting action will respond all hitting sequences, sorted by the descending order of sequence identify. Crosslinks to the genome browser and alignment results are also provided for the hits. Worthy of mention, the library is allowed to be fully downloaded in the *fasta* format for free academic use.

### The Library-based Method Can Generate Better Transcriptome

To evaluate the feasibility of the transcript library as an alternative reference to genome, we made the performance comparison of library-based, genome-based, and *de novo* method in transcriptome assembly, using two external amphioxus RNA-seq datasets as samples. These two external datasets were determined by different research groups, under different sequencing conditions, and with different sequencing qualities. The comparison was demonstrated on several aspects of read mapping ratio, number of unique genes (unigenes), number of bridges, N_50_, sequence completeness, fragmentation ratio, and estimated assembly score. The results were illustrated in **Figure 4**. In general, the Trinity *de novo* assembly exhibited higher reads mapping ratio and more unigenes than both genome-based and library-based methods. However, the Trinity method also yielded the smallest N_50_ value, comparatively more potential bridges, and higher fragmentary ratio at the same time (**Figure 4**). Comparatively, the library-based method exhibited the best sequence completeness, the lowest fragmentation ratio, and the high assembly score.

**Figure 4.**
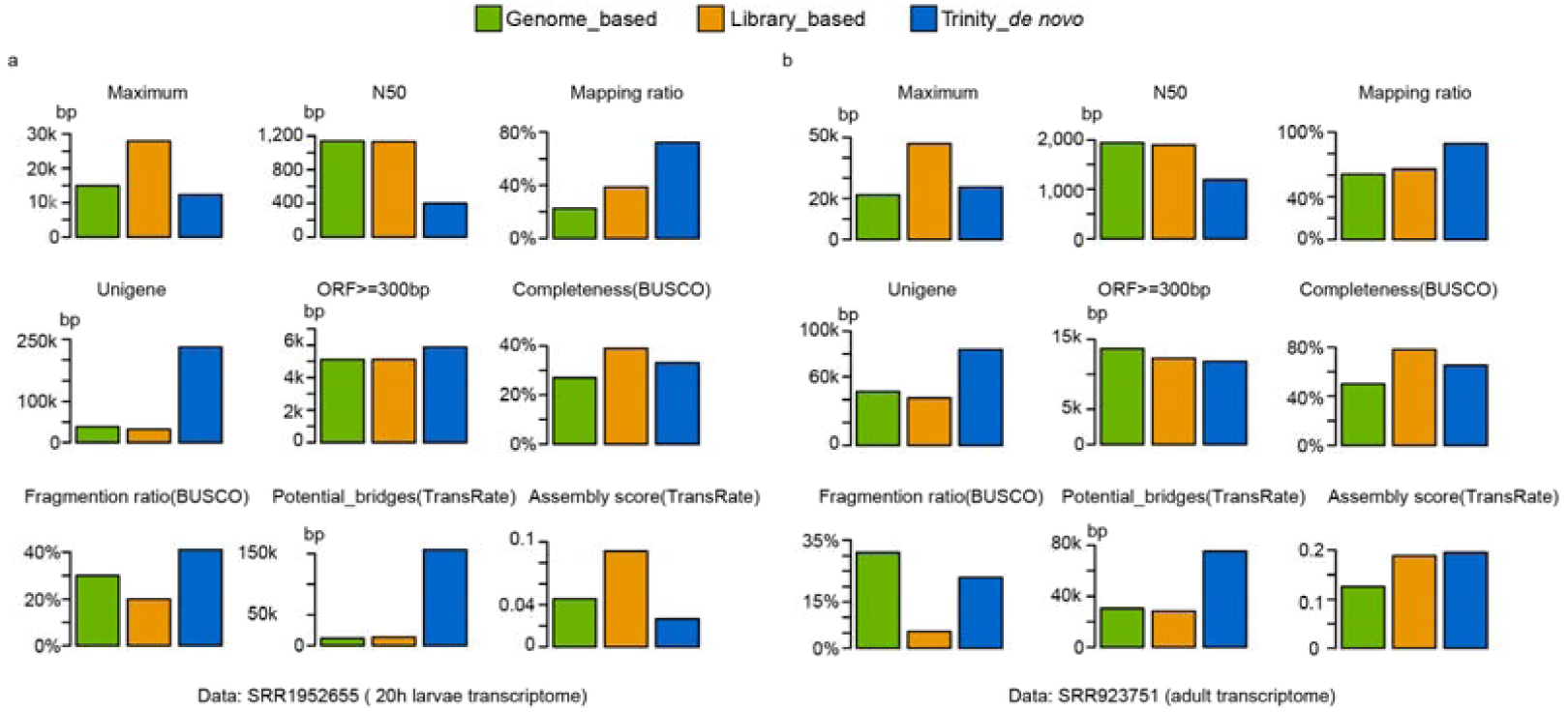
Performance comparison of transcriptome assembly using the library-based, genome-based, and *de novo* method.

In summary, the library-based method outperformed the genome-based method and the *de novo* method in transcriptome reconstruction. The integrative library allowed to utilize the sequencing reads to the largest extent in producing as more transcripts as possible; however at the same time, it overcome the fragmentary assembly problem in *de novo* method. Hence, the integrative transcript library can serve as an alternative reference to genome, especially under the circumstance when the genome is not good enough to yield a reliable and close-to-completeness transcriptome.

## Discussion

In this study, we proposed a novel computational method TransIntegrator, which can generate a library of full expressed transcripts for an organism in a genome-free manner via integrating multiple heterogeneous sequencing datasets. Taking the library as a reference, it is able to detect the “hidden” transcripts and gene isoforms in RNA-sequencing transcriptomes which are usually hard to be obtained by the conventional genome-based or *de novo* transcriptome assembly methods. Compared to the conventional methods such as HISAT, Trinity, or other combinational strategies, the library-based method introduced in this study has several merits: (1) The new method takes maximum advantages of NGS/TGS data heterogeneity to regenerate full landscape of transcript collection, regardless of low expression level and sequencing quality of single dataset. (2) The library can serve as an alternative reference in replace of genome for transcriptome assembly, which is consistently applicable for all transcriptome cases instead of single case. To our limited knowledge, it is a brand-new application. (3) The method is flexible and scalable that makes it easily implemented and rebuilt subject to user’s need and computing power. According to our experience, the *de novo* software IDBA-Trans used in this study can be easily replaced by Trinity or other state-of-art tools without substantially changing the workflow performance. (4) The method is economic and practical. Constructing a library of nearly full collection of transcripts just requires several RNA-seq datasets determined under different conditions. The completeness and diversity of the library can grow by incorporation of more heterogeneous transcriptome datasets. The more heterogeneous, the better.

Doubtlessly, the TransIntegrator method has its weakness. Its performance relies on the diversity of RNA-seq datasets. The *de novo* assembly and over-bridge will produce a portion of falsely assembled transcripts, especially for the genes with repetitive elements. These false transcripts could be partially excluded by several procedures such as reads mapping, transcript annotation, and longer sequencing reads. As the rapid widespread of sequencing technologies and the accumulation of RNA-seq datasets, the shortness will be overcome sooner and later. In particular, recent advances in TGS technology has brought the most potent to construct a complete transcript library.

Nevertheless, this study shows an example of using heterogeneous data maximally to overcome the gap of unsatisfied genome or no genome. It broadens the gene exploration from the static state to the dynamic state by providing a nearly complete transcript library as alternative reference to genome. Moreover, it brings systems biology research universally into the most organisms.

## Materials and Methods

### Sample preparation

The *Branchiostoma floridae* was originally introduced from Dr. Jr-Kai Yu’s laboratory (Institute of Cellular and Organismic Biology, Academia Sinica, Taiwan) and bred in our aquarium as described previously (Li, Yang, Shu, Chen, & Wang, 2012). Matured individuals were induced to spawn by heat shock method as described in previous study (Li, Shu, & Wang, 2013). Eggs released by three females were pooled together and then fertilized with sperm from one male. Embryos were raised in an incubator setting the temperature at 25 °C and the humidity at 85 % as previously described (X. Liu, Li, Feng, Yang, & Wang, 2013). Around 500 embryos at each of the following stages were collected and then lysed with one milliliter of Trizol reagent (Invitrogene Co.): egg, fertilized egg (ten minutes after fertilization), 2 cells, 32 cells, 256 cells, early gastrula, late gastrula, early neurula (just beginning to rotate), interim neurula (with five somite) and late neurula (onset of muscular movement). Total RNA were prepared according to the manual of Trizol reagent and then sent to Novogene Co. (China) for sequencing library construction and RNA-seq.

### The next generation sequencing (NGS) of the embryo RNA samples

The RNA library was constructed by the Novogene Co. (China). For each sample, a total amount of 3μg RNA was used as input material for library preparation. The polyadenylated (poly(A)) RNA was purified from total RNA using poly-T oligo-attached magnetic beads. Fragmentation was carried out using divalent cations under elevated temperature in NEBNext First Strand Synthesis Reaction Buffer (5X). After reverse transcription, the fragments were purified with AMPure XP system (Beckman Coulter, Beverly, USA), and the cDNA fragments with length in 150∼200bp were preferentially selected for PCR. The 3μl USER Enzyme (NEB, USA) was used for size-selected, adaptor-ligated cDNA at 37°C for 15 min followed by 5 min at 95 °C before PCR. Then PCR was performed with Phusion High-Fidelity DNA polymerase, Universal PCR primers and Index (X) Primer. At last, PCR products were purified (AMPure XP system) and library quality was assessed on the Agilent Bioanalyzer 2100 system. The RNA sequencing was demonstrated on the Illumina HiSeq™ 2000 system in a 125-base pair (bp) paired-end mode.

### The third-generation sequencing (TGS) of the embryo RNA samples and data processing

In this study, around 500 embryos were collected from each of several amphioxus embryo development stages, including egg, late gastrula, early neurula (3-som), middle neurula (8-som), late neurula, 3-branchial juvenile, and metamorphosis juvenile. The biological samples were mixed for total RNA extraction using the same protocol as described above. The total RNA samples were subsequently sent to the Novogene Co. (China) company for library construction and proceeded to the full-length cDNA library sequencing. The total amount of 3μg RNA was used for library construction and subsequently sequenced by single molecule real-time sequencing technology based on Pacbio rsII platform. The original sequencing data were treated by the official Pacbio software isoseq3 for standard processing.

### The computational workflow TransIntegrator for constructing the integrative transcript library

The computational workflow accepted the RNA-seq datasets *.*fq* and *.*fasta* file as input; and it output the transcript library file in *.*fasta* format. The workflow consisted of four major steps: (1) *De novo* assembly of NGS datasets, (2) Representative sequence search, (3) Re-assembly, and (4) Refinement and annotation (**Figure 1**).

#### RNA-seq data preprocessing

Before proceeding the NGS transcriptome assembly, a quality control on the raw sequencing data was made to exclude low-quality reads as the follow steps: 1) remove adaptor-contained reads; 2) Remove N-content > 10% reads; 3) Remove low quality reads that the percent of base with sQ <= 5 is more than 50% (of whole reads).

#### De novo assembly

In this study, we have tried implementing three frequently-used *de novo* methods Oases (Schulz et al., 2012), Trinity (Grabherr et al., 2011), and IDBA-Trans (Peng et al., 2013) into the workflow. Although the Trinity had several advantages in single transcriptome assembly, we still chose the IDBA-Trans (version 1.1.1) software for the initial *de novo* single transcriptome assembly since it took less computational time and was more sensitivity in detecting low-expressed transcripts (Peng et al., 2013). However, the Trinity is an option if the computational power is adequate. For every single transcriptome assembly by IDBA-Trans, a series of kmer size (from 90 to 125, step=5) were tried to elect the right kmer that gave the largest transcript number and the longest N50 (**Table S4**).

#### Representative sequence search

The representative sequence search was the critical step, in which the pre-assembled NGS datasets and TGS dataset were integrated. The *de novo* assembled NGS transcriptomes contain a large number of fragmentary sequences. To remove the identical or inclusive short fragments and at the same time keep those putative isoform fragments, we used CD-HIT-EST (version 4.5.4)(Y. Huang, Niu, Gao, Fu, & Li, 2010) to cluster all sequences and elect representatives. Two parameters were crucial for electing representative sequences: the sequence identity threshold (-c) and the alignment coverage (-aS). After several rounds of experiments, we determined the optimal parameters of identify threshold >90% and alignment coverage >80% for choosing representatives (**Figure S1**). Accordingly, we clustered sequences by similarity, discarded the shorter sequences, and elected the longest sequences as representatives. These representatives formed the pre-library of transcripts. For better computational efficiency, we started the data integration from the TGS dataset and the best quality RNA-seq datasets that have comparatively deep sequencing depth and long sequencing length. According to our experience, the two parameters (-c and -aS) could be adjusted subject to the estimated genetic variation level of the studying organisms. The large genetic variation expected comparatively smaller threshold value to retain isoforms.

#### Re-assembly

The pre-library of transcripts still contained a number of short sequences that were not subsets of others; however, they may possess overlapping fragments at either flank of sequences. These short sequences had chances to link together via the overlapping fragments to form longer sequences. Therefore, we used CAP3 (version 12/21/07)(X. Huang & Madan, 1999) to re-assemble sequences; any two sequences that had more than 40bp overlaps (>90% sequence identity) were bridged and extended to a new long sequence.

#### Refinement and library annotation

Lastly, we refined the library by removing sequences <300bp. After refinement, we achieved the integrative transcript library. We annotated the transcripts using the tool Annocript. The Annocript is a workflow to annotate transcriptome. Annotation mainly includes protein annotation, conservative domain annotation, open reading frame (ORF) prediction, go function annotation, enzyme pathway annotation, and ncRNA annotation (including the annotation of rRNA, tRNA, snRNA, snoRNA, miRNA, and lncRNA).

### The workflow running environment

The computational workflow TransIntegrator worked under the Linux operation system. It consisted of several component tools, including IDBA-Trans, CD-HIT and CAP3. Installation of these tools should follow the guidelines provided by the developers. Besides, the TransIntegrator was coded using Python language, which required the pre-installed Python environment. For user convenience, we packed all component tools together except Python into a software package; however, the licenses of these tools still belong to their developers. The workflow allowed modifying several key parameters for optimal performance of component tools. We have deposited the TransIntegrator package at https://github.com/BADDxmu/IncDNA.git and http://www.bio-add.org/InTrans/ for free downloading.

### Reference-based transcriptome assembly and annotation

We adopted the TopHat (version 2.0.11) (Trapnell, Pachter, & Salzberg, 2009) for reference-based transcriptome assembly. The reference could be either the amphioxus genome (JGIv2.0) or the integrative transcript library built in this study.

### Performance comparison of different transcriptome assembly methods

We compared the library-based method, the genome-based method, and *de novo* Trinity method to evaluate their performance on single transcriptome assembly using two external amphioxus RNA-seq datasets from public resources (SRA: SRR923751 and SRR1952655) as the samples. These two datasets were determined under different experiment conditions and had different sequencing quality. In the comparison, we used TopHat (version 2.0.11) (Trapnell et al., 2009)+Cufflink (version 2.0.11) (Ghosh & Chan, 2016) pipeline for the reference-based (refer to both library and genome) transcriptome assembly. The comparison adopted several statistical parameters like reads mapping ratio, N_50_, and unique gene (unigene) number to evaluate the quality of transcriptome assembly (Moreton et al., 2015). We also measured the completeness and fragmentation ratio of transcripts using the tool BUSCO (Simao, Waterhouse, Ioannidis, Kriventseva, & Zdobnov, 2015), and estimated the potential bridge and assembly score by TransRate (Smith-Unna, Boursnell, Patro, Hibberd, & Kelly, 2016).

## Acknowledgements

Financial support from the Natural Science Foundation of China (NSFC#31671362 and 31271405) is gratefully acknowledged.

## Figure legends

**Figure S1.**
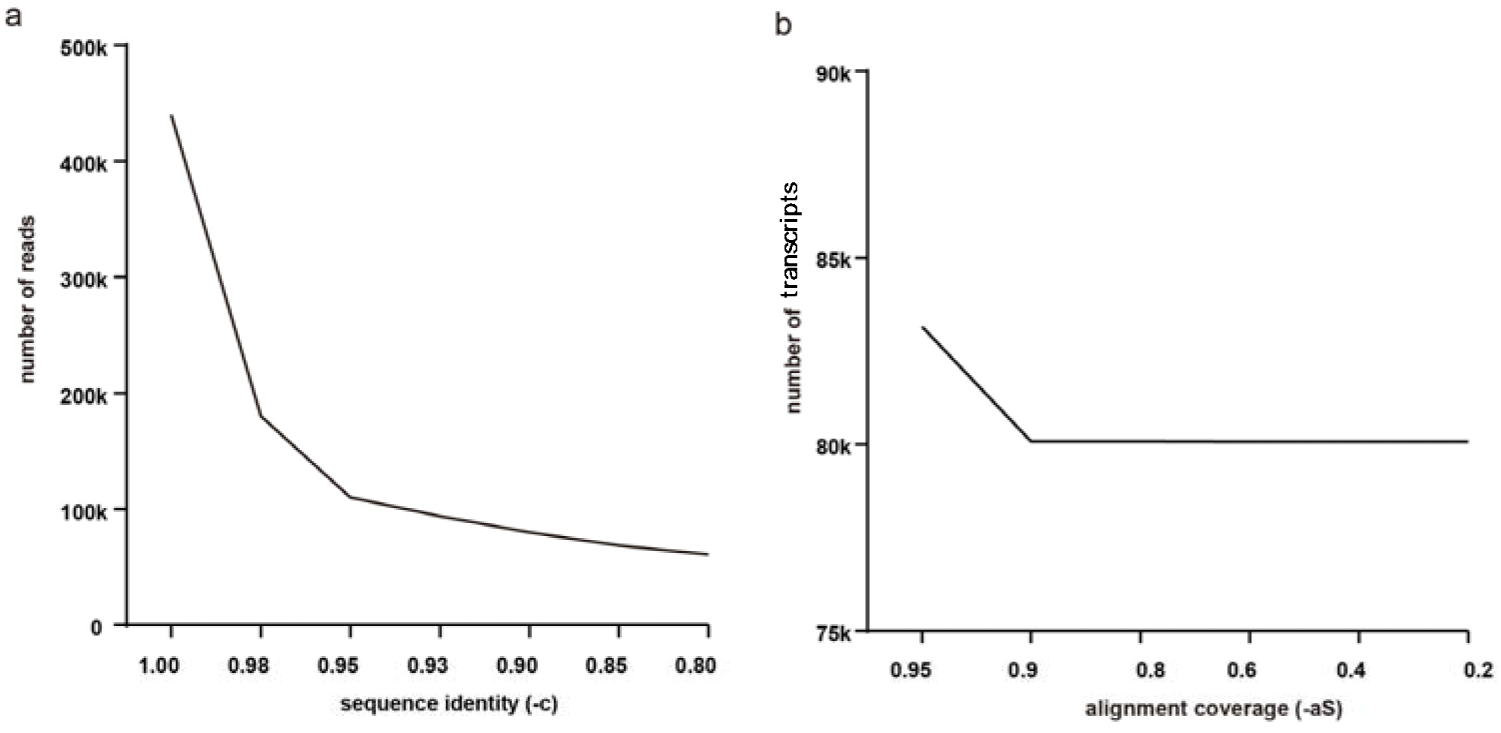
Determination of CD-HIT-EST parameters for transcript library integration. (a) Integration of transcript library reads by different sequence identity (-c). The number of reads drops sharply along with the decrease of sequence identity until -c=0.95 and thereafter. (b) Integration of transcript library under different alignment coverages (-aS) and constant sequence identity (-c)=0.9. The transcripts number drops quickly with the decrease of -aS and remains constant of about 80,000 sequences after -aS=0.9.

## References

Armero, A., Baudouin, L., Bocs, S., & This, D. (2017). Improving transcriptome de novo assembly by using a reference genome of a related species: Translational genomics from oil palm to coconut. PLoS One, 12(3), e0173300. doi: 10.1371/journal.pone.0173300

Bao, Y., Xu, S., Jing, X., Meng, L., & Qin, Z. (2015). De novo assembly and characterization of Oryza officinalis leaf transcriptome by using RNA-seq. BioMed Research International, 2015, 982065. doi: 10.1155/2015/982065

Benito-Gutiérrez, È. (2011). Amphioxus as a Model for Mechanisms in Vertebrate Development: John Wiley & Sons, Ltd.

Bouyioukos, C., Moscou, M. J., Champouret, N., Hernández-Pinzón, I., Ward, E. R., & Wulff, B. B. H. (2013). Characterisation and Analysis of the Aegilops sharonensis Transcriptome, a Wild Relative of Wheat in the Sitopsis Section. PLOS ONE, 8(8), e72782. doi: 10.1371/journal.pone.0072782

Buels, R., Yao, E., Diesh, C. M., Hayes, R. D., & Holmes, I. H. (2016). JBrowse: A dynamic web platform for genome visualization and analysis. Genome Biology, 17(1).

Cerveau, N., & Jackson, D. J. (2016). Combining independent de novo assemblies optimizes the coding transcriptome for nonconventional model eukaryotic organisms. BMC Bioinformatics, 17(1), 525. doi: 10.1186/s12859-016-1406-x

Chaisson, M. J., Wilson, R. K., & Eichler, E. E. (2015). Genetic variation and the de novo assembly of human genomes. Nat Rev Genet, 16(11), 627–640. doi: 10.1038/nrg3933

Chen, C. C., Lin, W. D., Chang, Y. J., Chen, C. L., & Ho, J. M. (2012). Enhancing de novo transcriptome assembly by incorporating multiple overlap sizes. ISRN Bioinform, 2012, 816402. doi: 10.5402/2012/816402

Chen, G., Yin, K., Wang, C., & Shi, T. (2011). De novo transcriptome assembly of RNA-Seq reads with different strategies. Sci China Life Sci, 54(12), 1129–1133. doi: 10.1007/s11427-011-4256-9

Gan, X., Stegle, O., Behr, J., Steffen, J. G., Drewe, P., Hildebrand, K. L., Mott, R. (2011). Multiple reference genomes and transcriptomes for Arabidopsis thaliana. Nature, 477(7365), 419–423.

Ghosh, S., & Chan, C. K. (2016). Analysis of RNA-Seq Data Using TopHat and Cufflinks. Methods Mol Biol, 1374, 339–361. doi: 10.1007/978-1-4939-3167-5_18

Grabherr, M. G., Haas, B. J., Yassour, M., Levin, J. Z., Thompson, D. A., Amit, I., Regev, A. (2011). Full-length transcriptome assembly from RNA-Seq data without a reference genome. Nat Biotechnol, 29(7), 644–652. doi: 10.1038/nbt.1883

Haas, B. J., Papanicolaou, A., Yassour, M., Grabherr, M., Blood, P. D., Bowden, J., Regev, A. (2013). De novo transcript sequence reconstruction from RNA-seq using the Trinity platform for reference generation and analysis. Nat Protoc, 8(8), 1494–1512. doi: 10.1038/nprot.2013.084

Hu, Z., Scott, H. S., Qin, G., Zheng, G., Chu, X., Xie, L., Wei, C. (2015). Revealing Missing Human Protein Isoforms Based on Ab Initio Prediction, RNA-seq and Proteomics. Sci Rep, 5, 10940. doi: 10.1038/srep10940

Huang, S., Chen, Z., Yan, X., Yu, T., Huang, G., Yan, Q., Xu, A. (2014). Decelerated genome evolution in modern vertebrates revealed by analysis of multiple lancelet genomes. Nat Commun, 5, 5896. doi: 10.1038/ncomms6896

Huang, X., Chen, X. G., & Armbruster, P. A. (2016). Comparative performance of transcriptome assembly methods for non-model organisms. BMC Genomics, 17, 523. doi: 10.1186/s12864-016-2923-8

Huang, X., & Madan, A. (1999). CAP3: A DNA sequence assembly program. Genome Res, 9(9), 868–877. doi: 10.1101/gr.9.9.868

Huang, Y., Niu, B., Gao, Y., Fu, L., & Li, W. (2010). CD-HIT Suite: a web server for clustering and comparing biological sequences. Bioinformatics, 26(5), 680–682. doi: 10.1093/bioinformatics/btq003

Kim, D., Langmead, B., & Salzberg, S. L. HISAT: a fast spliced aligner with low memory requirements. Nature Methods, 12(4), 357–360.

Korlach, J. (2013). Understanding Accuracy in SMRT Sequencing.

Lowe, R., Shirley, N., Bleackley, M., Dolan, S., & Shafee, T. (2017). Transcriptomics technologies. PLoS Comput Biol, 13(5), e1005457. doi: 10.1371/journal.pcbi.1005457

Marlétaz, F., Firbas, P. N., Maeso, I., Tena, J. J., Bogdanovic, O., Perry, M., Irimia, M. (2018). Amphioxus functional genomics and the origins of vertebrate gene regulation. Nature, 564(7734), 64–70. doi: 10.1038/s41586-018-0734-6

Moreton, J., Izquierdo, A., & Emes, R. D. (2015). Assembly, Assessment, and Availability of De novo Generated Eukaryotic Transcriptomes. Frontiers in Genetics, 6, 361. doi: 10.3389/fgene.2015.00361

Nakasugi, K., Crowhurst, R., Bally, J., & Waterhouse, P. (2014). Combining transcriptome assemblies from multiple de novo assemblers in the allo-tetraploid plant Nicotiana benthamiana. PLoS One, 9(3), e91776. doi: 10.1371/journal.pone.0091776

Orgeur, M., Martens, M., Borno, S. T., Timmermann, B., Duprez, D., & Stricker, S. (2018). A dual transcript-discovery approach to improve the delimitation of gene features from RNA-seq data in the chicken model. Biol Open, 7(1). doi: 10.1242/bio.028498

Oulion, S., Bertrand, S., Belgacem, M. R., Le Petillon, Y., & Escriva, H. (2012). Sequencing and analysis of the Mediterranean amphioxus (Branchiostoma lanceolatum) transcriptome. PLoS One, 7(5), e36554. doi: 10.1371/journal.pone.0036554

Peng, Y., Leung, H. C., Yiu, S. M., Lv, M. J., Zhu, X. G., & Chin, F. Y. (2013). IDBA-tran: a more robust de novo de Bruijn graph assembler for transcriptomes with uneven expression levels. Bioinformatics, 29(13), i326–334. doi: 10.1093/bioinformatics/btt219

Peng, Y., Leung, H. C. M., Yiu, S. M., & Chin, F. Y. L. (2010). IDBA A Practical Iterative de Bruijn Graph De Novo Assembler. Research in Computational Molecular Biology, Proceedings, 6044, 426–440.

Pertea, M., Kim, D., Pertea, G. M., Leek, J. T., & Salzberg, S. L. (2016). Transcript-level expression analysis of RNA-seq experiments with HISAT, StringTie and Ballgown. Nat Protoc, 11(9), 1650–1667. doi: 10.1038/nprot.2016.095

Putnam, N. H., Butts, T., Ferrier, D. E., Furlong, R. F., Hellsten, U., Kawashima, T., Rokhsar, D. S. (2008). The amphioxus genome and the evolution of the chordate karyotype. Nature, 453(7198), 1064–1071. doi: 10.1038/nature06967

Putnam, N. H., Butts, T., Ferrier, D. E. K., Furlong, R. F., Hellsten, U., & et al. The amphioxus genome and the evolution of the chordate karyotype. Nature, 453(7198), 1064-1071,qt0003.

Rupp, O., Becker, J., Brinkrolf, K., Timmermann, C., Borth, N., Puhler, A., Goesmann, A. (2014). Construction of a public CHO cell line transcript database using versatile bioinformatics analysis pipelines. PLoS One, 9(1), e85568. doi: 10.1371/journal.pone.0085568

Schulz, M. H., Zerbino, D. R., Vingron, M., & Birney, E. (2012). Oases: robust de novo RNA-seq assembly across the dynamic range of expression levels. Bioinformatics, 28(8), 1086–1092. doi: 10.1093/bioinformatics/bts094

Sharon, D., Tilgner, H., Grubert, F., & Snyder, M. (2013). A single-molecule long-read survey of the human transcriptome. Nat Biotechnol, 31(11), 1009–1014. doi: 10.1038/nbt.2705

Simao, F. A., Waterhouse, R. M., Ioannidis, P., Kriventseva, E. V., & Zdobnov, E. M. (2015). BUSCO: assessing genome assembly and annotation completeness with single-copy orthologs. Bioinformatics, 31(19), 3210–3212. doi: 10.1093/bioinformatics/btv351

Smith-Unna, R., Boursnell, C., Patro, R., Hibberd, J. M., & Kelly, S. (2016). TransRate: reference-free quality assessment of de novo transcriptome assemblies. Genome Res, 26(8), 1134–1144. doi: 10.1101/gr.196469.115

Sohn, J. I., & Nam, J. W. (2016). The present and future of de novo whole-genome assembly. Brief Bioinform. doi: 10.1093/bib/bbw096

Surget-Groba, Y., & Montoya-Burgos, J. I. (2010). Optimization of de novo transcriptome assembly from next-generation sequencing data. Genome Res, 20(10), 1432–1440. doi: 10.1101/gr.103846.109

Trapnell, C., Pachter, L., & Salzberg, S. L. TopHat: discovering splice junctions with RNA-Seq. Bioinformatics, 25(9), 1105–1111.

Trapnell, C., Pachter, L., & Salzberg, S. L. (2009). TopHat: discovering splice junctions with RNA-Seq. Bioinformatics, 25(9), 1105–1111. doi: 10.1093/bioinformatics/btp120

Trapnell, C., Roberts, A., Goff, L., Pertea, G., Kim, D., Kelley, D. R., Pachter, L. Differential gene and transcript expression analysis of RNA-seq experiments with TopHat and Cufflinks. NATURE PROTOCOLS, 7(3), 562–578.

Wu, P., Zhang, H., Lin, W., Hao, Y., Ren, L., Zhang, C., He, F. (2014). Discovery of novel genes and gene isoforms by integrating transcriptomic and proteomic profiling from mouse liver. J Proteome Res, 13(5), 2409–2419. doi: 10.1021/pr4012206

Yang, Y., & Smith, S. A. (2013). Optimizing de novo assembly of short-read RNA-seq data for phylogenomics. BMC Genomics, 14, 328. doi: 10.1186/1471-2164-14-328

Zhao, Q. Y., Wang, Y., Kong, Y. M., Luo, D., Li, X., & Hao, P. (2011). Optimizing de novo transcriptome assembly from short-read RNA-Seq data: a comparative study. BMC Bioinformatics, 12 Suppl 14, S2. doi: 10.1186/1471-2105-12-S14-S2

