## Supplemental table1-4 for "Integrate Heterogeneous NGS and TGS Data to Boost Genome-free Transcriptome Research"

**Table S1. The genome information of amphioxus subspecies**

| Organisms | Sequencing system | Sequencing Depth | Genome size | Gene number | Protein number |
| --- | --- | --- | --- | --- | --- |
| *B. floridae* | Sanger | 17 | 522Mb | 21,954 | 50,817 |
| *B. belcheri* | Roche454/Illumina | 30/70 | 426Mb | 26,706 | 34,675 |
| *B. lanceolatum* | Illumina | 150 | 495Mb | 76,831 | 38,739 |

**Table S2. The sequencing information of transcriptomes determined at ten Amphioxus embryo development stages.**

| Samples | Clean reads | Clean  bases | Q20(%) | Q30(%) | GC(%) |
| --- | --- | --- | --- | --- | --- |
| egg_1 | 34,524,985 | 4.32G | 97.21 | 94.18 | 47.26 |
| egg_2 | 34,524,985 | 4.32G | 95.61 | 91.62 | 47.24 |
| fertilized egg_1 | 26,533,480 | 3.32G | 96.93 | 93.63 | 45.49 |
| fertilized egg_2 | 26,533,480 | 3.32G | 93.49 | 88.10 | 45.40 |
| 2 cells_1 | 33,476,307 | 4.18G | 96.83 | 93.44 | 45.21 |
| 2 cells_2 | 33,476,307 | 4.18G | 93.50 | 88.15 | 45.10 |
| 32 cells_1 | 30,354,192 | 3.79G | 96.83 | 93.41 | 47.07 |
| 32 cells_2 | 30,354,192 | 3.79G | 93.43 | 88.03 | 46.96 |
| 256 cells_1 | 27,005,360 | 3.38G | 96.82 | 93.41 | 47.50 |
| 256 cells_2 | 27,005,360 | 3.38G | 92.37 | 86.36 | 47.32 |
| early gastrulae_1 | 28,511,401 | 3.56G | 96.87 | 93.50 | 47.16 |
| early gastrulae_2 | 28,511,401 | 3.56G | 93.69 | 88.41 | 47.07 |
| late gastrulae_1 | 27,281,807 | 3.41G | 97.42 | 94.55 | 47.92 |
| late gastrulae_2 | 27,281,807 | 3.41G | 95.64 | 91.64 | 47.89 |
| early neurula_1 | 22,818,963 | 2.85G | 96.64 | 93.04 | 49.63 |
| early neurula_2 | 22,818,963 | 2.85G | 92.68 | 86.72 | 49.49 |
| interim neurula_1 | 36,249,400 | 4.53G | 99.02 | 97.22 | 47.86 |
| interim neurula_2 | 36,249,400 | 4.53G | 97.49 | 94.64 | 47.94 |
| late neurula_1 | 39,939,409 | 4.99G | 99.00 | 97.18 | 48.09 |
| late neurula_2 | 39,939,409 | 4.99G | 97.36 | 94.41 | 48.17 |

**Table S3. The raw data of single molecule real-time sequencing**

| samples | Subreads(G) | subreads | Average length(bp) | N50 |
| --- | --- | --- | --- | --- |
| *7-Stage mixed RNA samples* | 15.74 | 5,502,217 | 2,861 | 3,208 |

**Table S4. *De novo* transcriptome assembly under different kmer using RNA-seq data at the “egg” stage as example**

| Kmer | Contigs | N50(bp) | Maximum(bp)^*^ | Mean(bp) | Total(bp) |
| --- | --- | --- | --- | --- | --- |
| 90 | 43,817 | 2,332 | 15,649 | 1,339 | 58,671,285 |
| 95 | 45,411 | 2,602 | 16,533 | 1,490 | 67,665,781 |
| 100 | 44,257 | 2,690 | 16,533 | 1,555 | 68,840,174 |
| 105 | 43,985 | 2,712 | 16,533 | 1,586 | 69,774,912 |
| 110 | 43,649 | 2,774 | 16,533 | 1,616 | 70,560,477 |
| 115 | 44,085 | 2,830 | 16,533 | 1,641 | 72,350,603 |
| 120 | 45,100 | 2,848 | 16,533 | 1,652 | 74,523,193 |

* maximum length of contig in base pair (bp).
